# Testing The Protective Function of Flash Coloration Using Wild Avian Predators

**DOI:** 10.64898/2026.10.15.759298

**Authors:** Theo Brown, Sandra Winters, Pierre d’Halluin, George RA Hancock, Kelly Kittus, Toomas Tammaru, Anna Riu, Johanna Mappes

## Abstract

Many prey reveal conspicuous color patches during movement that become concealed when they stop. Such abrupt changes in appearance have been proposed to disrupt predator tracking and make prey more difficult to relocate, providing the basis of the flash coloration hypothesis. However, empirical support remains inconsistent and comes largely from experiments using human observers or simplified artificial prey. Here, we used touchscreen-equipped operant chambers to test the flash coloration hypothesis with wild blue tits (*Cyanistes caeruleus*) searching for moving digital prey modelled on Catocala moths. The digital prey revealed either conspicuous or cryptic hindwings during flight before adopting a cryptic resting appearance. Contrary to predictions, conspicuous hindwings neither increased prey survival nor delayed prey relocation compared to moths with cryptic hindwings. However, conspicuous hindwings also did not incur detectable predation costs. In contrast, background strongly affected prey detection, with moths surviving better on realistic bark than on smooth grey backgrounds. Our results provide no evidence that conspicuous colors revealed during movement enhance prey survival in this context, suggesting that any protective effect attributed to flash coloration may be weak or strongly dependent on ecological conditions.

**Lay Summary:** Many animals reveal bright colors while moving but conceal them when stationary. This sudden change in appearance has been proposed to confuse predators and make prey harder to relocate. We tested this idea using wild blue tits hunting digital moths. Bright hindwings revealed during flight did not increase moth survival or delay relocation, but they also carried no detectable predation cost. The protective value of these movement-dependent color changes may therefore be weak or depend strongly on ecological context.

## Introduction

Many prey species undergo striking changes in appearance when they move. Conspicuous color patches that are hidden at rest may become exposed during movement and then disappear again when the prey stops (Stevens and Cuthill 2006; Stevens and Merilaita 2009; Caro and Koneru 2021; Scott-Samuel et al. 2023). Two commonly cited examples are the blue-winged grasshopper (*Oedipoda caerulescens*) and *Catocala* moths, whose brightly colored hindwings are visible during flight but concealed beneath cryptic forewings at rest (Sargent 1978; Ingalls 1993; Kim et al. 2020). Such movement-dependent changes in appearance have long been proposed to reduce predation by confusing or misdirecting predators pursuing moving prey (Cott 1940). Testing this proposed anti-predator function is experimentally challenging, however, because controlled experiments must simultaneously recreate prey movement, rapid changes in appearance, and predator tracking behavior (Stevens and Cuthill 2006; Stevens and Merilaita 2009; Cuthill et al. 2017; Ruxton et al. 2018; Murali and Kodandaramaiah 2020; Caro and Koneru 2021; Scott-Samuel et al. 2023).

Cott (1940) originally coined the term “flash coloration” to describe both the form and putative function of these color patches. However, terminology surrounding flash coloration remains ambiguous, and terms such as flash coloration, flash behaviors, and flash displays have all been used to describe similar phenomena (Loeffler-Henry et al. 2018; Murali 2018; Bae et al. 2019; Loeffler-Henry et al. 2021; No et al. 2024 Sept 20; Vieira Silva et al. 2024). Here, we use *flash coloration* to refer specifically to conspicuous color patches that are visible during motion but concealed when the animal is stationary (i.e., the form of the color pattern) and the *flash coloration hypothesis* to describe the proposed function of flash colors. The hypothesis predicts that the abrupt transition from conspicuous moving and cryptic stationary appearances disrupts predator tracking or prey relocation, thereby increasing prey survival. Mechanistically, this transition may disrupt the continuity of the predator’s visual representation of the prey, requiring the predator to update both its search image and the visual features used for target identification, potentially preventing or delaying rediscovery (Langley 1996; Bond and Kamil 1999; Loeffler-Henry et al. 2018).

Empirical support for the flash coloration hypothesis remains inconsistent (Table 1). Some studies have reported increased prey survival or disrupted predator tracking associated with movement-dependent conspicuous signals (Loeffler-Henry et al. 2018; Bae et al. 2019; No et al. 2024 Sept 20; Silvasti et al. 2024) whereas others have found weak or absent effects (Murali 2018; Potter 2023; Vieira Silva et al. 2024). Existing experiments also differ substantially in predator identity, prey realism, experimental design, and the metrics used to quantify predator responses. Most previous work has relied on human participants, domestic chicks, or simplified artificial prey such as moving colored squares (See Table 1). Although such systems allow strong experimental control, they may fail to capture important aspects of natural predator-prey interactions. Consequently, it remains unclear whether concealed conspicuous color patches genuinely disrupt the tracking capacity of wild, natural predators. Interactive touchscreen systems offer a way to address some of these limitations by combining precisely controlled dynamic prey with wild natural predators under repeatable experimental conditions.

**Table 1.** Summary of published experimental studies testing the flash coloration hypothesis. The table lists predator species, type of flash coloration used, experimental system, and main results. The evidence to date remains mixed across systems and taxa.

| Prey<br>Appearance | Predator<br>Species | System | Flash color –<br>cryptic color | Findings | Clear evidence of<br>conspicuous<br>colors being the<br>most effective<br>defense | Reference |
| --- | --- | --- | --- | --- | --- | --- |
| Digital Squares | Human | Screen-based<br>game | Red - Green | 20% increased survival rate | Yes | Loeffler-Henry<br>et al. 2018 |
| Digital Squares | Human | Screen-based<br>game | White –<br>Black/Grey | Flash colors were as<br>effective as continuous<br>grey | No | Murali 2018 |
| Digital Squares | Human | Screen-based<br>game | Red - Green | Flash effectiveness<br>dependent on large<br>stimulus size | Yes | Bae et al. 2019 |
| Digital Squares | Human | Screen-based<br>game | Red/Orange/Y<br>ellow –<br>Green/Grey | Flash coloration is more<br>effective when color is<br>aposematic | Yes | Kim et al. 2020 |
| Digital Squares | Human | Screen-based<br>game | Red –<br>Green/Beige | Flash effectiveness<br>dependent on FID | Yes | Loeffler-Henry<br>et al. 2021 |
| Digital Frogs | Chicks | Projector | Blue – Black<br>and White | No survival advantage v.<br>continuous gray | No | Potter 2023 |
| Digital Squares | Chicks | Projector | Red/Brown -<br>Green | 22% increased survival but<br>brown flash color as<br>effective as red | No | No et al. 2024 |
| Morpho Butterfly | Birds | Capture-mark-<br>recapture | Blue - Brown | No survival advantage from<br>flash color patch | No | Vieira-Silva et<br>al. 2024 |

Moths of the genus Catocala provide a particularly suitable system for testing the flash coloration hypothesis. These large nocturnal moths typically rest on tree trunks, where their cryptic forewings closely resemble bark, but expose brightly colored hindwings during flight before concealing them again upon landing. Catocala moths have therefore long been regarded as classic examples of flash coloration (Sargent 1978; Ingalls 1993), yet the hypothesis has not been experimentally tested in this iconic system. Birds are important predators of adult Lepidoptera, making avian responses particularly relevant for evaluating the proposed anti-predator function of these concealed conspicuous displays.

Here, we used touchscreen-equipped operant chambers (TOCs) to test whether concealed conspicuous hindwing coloration influences predator search behavior and prey survival using wild blue tits (*Cyanistes caeruleus*) as predators. Blue tits are well-established model predators in studies of avian vision and moth coloration and have been shown to predate adult lepidoptera (Betts 1955; Hart et al. 2000; Rönkä et al. 2018; Burdfield Steel et al. 2025). We presented birds with digital moths modelled on *Catocala* species, possessing either conspicuous or cryptic hindwings during flight before the moth landing in the same cryptic resting posture. We predicted that conspicuous hindwings would make prey more difficult to relocate after landing, resulting in longer search times and increased survival relative to moths with cryptic hindwings. Trials were conducted on tree bark or a solid grey background, allowing us additionally to test the prediction that resting prey would be more difficult to relocate on a complex background than on a uniform background (Merilaita 2003; Dimitrova and Merilaita 2010; Xiao and Cuthill 2016). By combining controlled dynamic prey presentations with ecologically relevant prey signals and wild predators, our study provides a direct experimental test of the flash coloration hypothesis.

## Methods

### Predators

Experiments were carried out at Lammi Biological Station, Finland (61°03’13.8"N 25°02’25.5"E) across two non-breeding seasons (Feb-Mar & Sept-Dec 2023). Wild blue tits were used as predators given their ability to learn complex foraging tasks, their rapid adjustment to captivity, and their status as generalist predators of insects (Betts 1955; Hämäläinen et al. 2017; Silvasti et al. 2024). All birds were experimentally naïve. Birds were kept in individual plywood cages on an 11 h:13 h (light:dark) schedule, and provided with food: peeled sunflower seeds, peanuts, and crushed suet and fresh water *ad libitum*. Food was removed approximately 90 minutes before training or experimental sessions began. The purpose of food deprivation was to increase motivation and to help overcome initial neophobia to new training stages. Training of each bird took place between 8:30 and 17:00 each day, after which the food was returned to the birds. Food intake across training and testing was carefully monitored to ensure sufficient food was provided. After completing the experiment, birds were aged, ringed, and released. Birds were not sexed given the relative difficulty in distinguishing male and female blue tits. Birds were kept captive in line with the Central Finland Regional Environment Centre Permit (VARELY/6870/2022).

### Experimental apparatus

We ran trials using touchscreen-equipped operant chambers (TOCs), which are controlled experimental apparatuses consisting of a large wooden plywood box (90 cm width x 90 cm height x 80 cm depth) which contains a computer monitor on one inside wall (Figure 1A).

**Figure 1.**
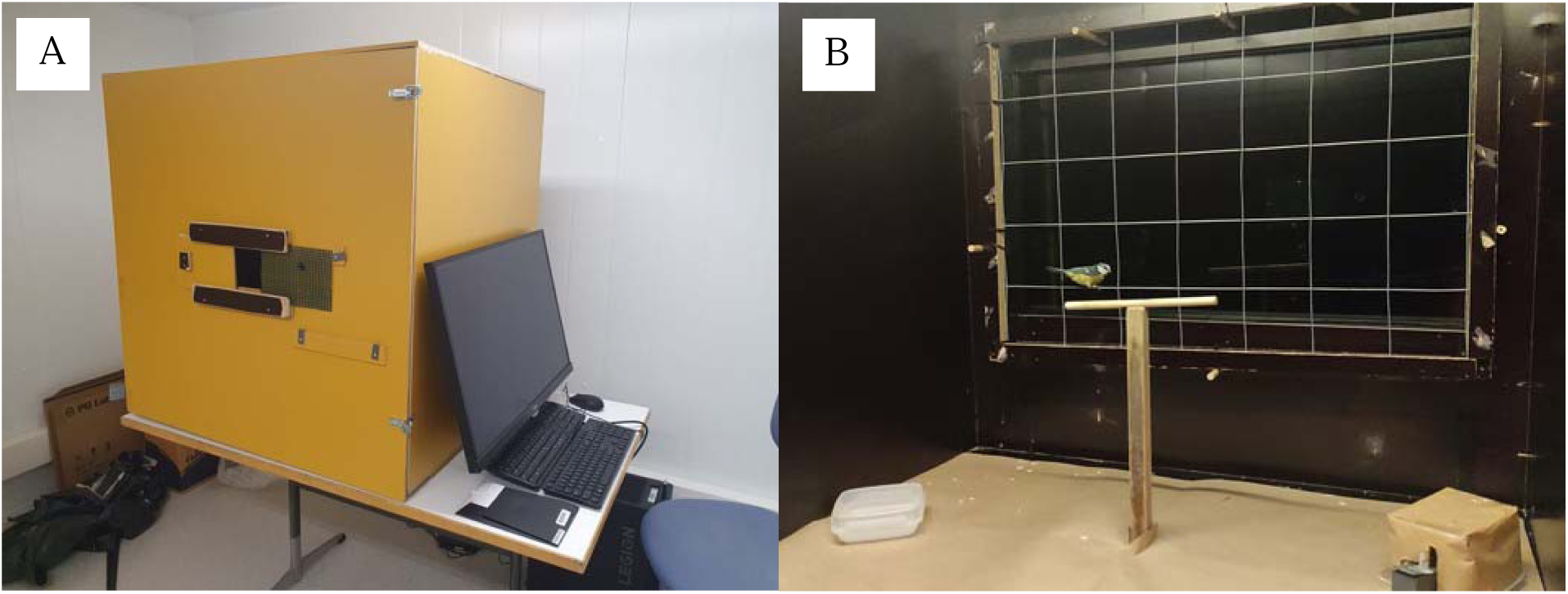
A. The exterior of the touchscreen-equipped operant chamber (TOC). There is a viewing slot in the front panel of the TOC and an external monitor for controlling the touchscreen. B. The inside of the TOC, with a bird on the perch. A metal grid overlays the monitor to allow the bird access to the whole screen. The remote-controlled food reward dispenser is on the bottom right.

The monitor is overlayed first by a 3.5mm thick acrylic sheet to protect the screen from scratches and then by an infrared frame (32” GreenTouch Multi-Points Infrared Touch Overlays) which detected pecks, allowing the monitor to function as touchscreen. In front of the monitor was a wire grid which allowed the bird to access any area of the screen. In the center of the box was a perch and the box was illuminated by an LED strip light on the ceiling. To ensure broad color representation with a fast refresh rate, high quality monitors were used (Republic of Gamers Swift PG329Q 32’’: 165 Hz, >99% of sRGB color space, 2560 x 1440 pixel resolution) and were paired with suitably powerful computers (Lenovo Legion T5 Ryzen 7 with a NVIDIA GeForce RTX 3070 GPU). For more details on the TOC apparatus, see Winters et al. 2026.

### Training procedures

We used a scaffolding approach to train birds to peck the digital moths on the monitor. Birds progressed from pecking paper moths with seeds glued to them, to pecking paper moths and receiving a seed from the remote-controlled box, and then to pecking digital moths against a grey background (RGB (uint8) = 181, 181, 181) on the screen. Once birds had successfully learned to peck static digital moths, they were then trained to interact with resting moths that glided across the screen (i.e., wings did not flap and hindwings were never revealed): birds had to peck a digital moth, which then moved a short distance before stopping, and then re-peck the same moth to receive a reward. To overcome any neophobia related to pecking moths on textured background images, birds were then trained to peck moths on a bark background by using similar images to those used in the experiment trials. The color of the moths used in training was a light red (RGB (uint8) = 196, 88, 86) and a light orange (RGB (uint8) = 177, 131, 81) color, chosen because they are approximately equidistant from the conspicuous (red and yellow) and grey hindwing colors used in the experimental stimuli (see below). This was to avoid biasing the birds towards any experimental stimuli during the training phase. See supplementary materials for full training details.

### Experimental design

Experimental trials involved birds pecking digital moths that moved dynamically across stationary background images. Each trial consisted of three stages: (1) the bird initiated the trial by pecking a visible resting moth positioned near the edge of the screen, (2) the moth performed a short (2.3s) random flight during which the hindwing colors were visible, and (3) after landing, the moth became cryptic and the bird searched for and attempted to relocate it on the background image. The experiment therefore specifically tested whether conspicuous hindwing displays shown during movement impaired a predator’s ability to relocate prey after landing.

Twenty-one wild blue tits, none of which had prior experience with the apparatus, each completed 18 trials in the experiment. Each bird was exposed to digital moths with three different hindwing colors (red, yellow, and grey; see Figure 2) and two background types (bark and smooth), resulting in six experimental conditions. The bark backgrounds were photographs of tree bark on which stationary moths appeared well camouflaged. The smooth backgrounds were generated by taking the average color of each bark image upon which the stationary moths were easy to detect. Using both a natural bark background and a smooth control background of equivalent average color allowed us to isolate the effect of natural background patterning while controlling for overall background luminance and color. Each condition was repeated three times per bird in a randomized block design, resulting in 18 trials per bird.

**Figure 2.**
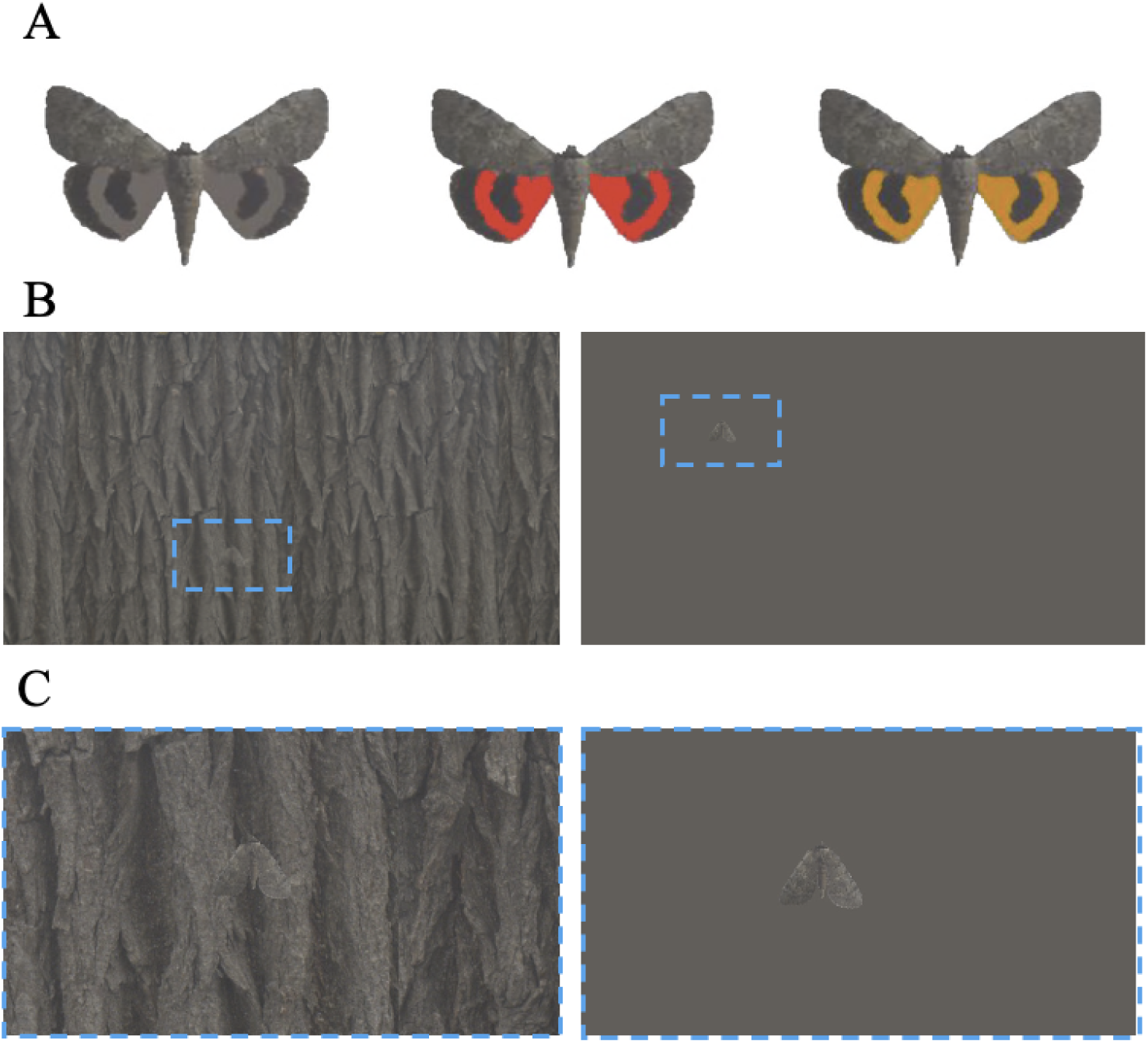
A. The three experimental stimuli. The cryptic (grey), red, and yellow hindwings are only visible when the moth is flying. B. Example images of the bark and smooth background types with a moth in resting posture. The outlined regions denote the moth locations. C. Zoomed images showing the moths on the bark and smooth background. Note that the resting moth is more conspicuous on the smooth background than on the bark background.

Each trial was initiated with a start screen containing a background image with a 6cm grey (RGB (uint8) = 181, 181, 181) border. A static 3cm x 2.5cm moth in the resting posture was randomly placed on the grey border. To start the trial, the bird had to peck the static moth. Pecking a clearly visible moth meant the bird was motivated to start the trial. After this first peck on the moth, the grey border was removed so the background image covered the entire screen for the rest of the trial and the moth started a 2.3 second moth flight at a speed of 5.6cm/s. During the flight, the moth would display one of the three hindwing color morphs before coming to rest on the background (see supplementary materials for full trial details and an example video of a trial).

Once the moth landed, the time taken for the bird to re-find the moth was recorded automatically by the computer and the time the bird spent actively looking for the moth was recorded by an observer using a stopwatch. The bird was deemed to be searching for the moth if it was either on the grid next to the screen or if it was on the perch and facing in the direction of the screen. Birds were allowed either two minutes of active searching or five minutes of total time to find the moth; if either of these criteria were met, the trial was concluded and the bird did not get a reward. Video footage of each trial was taken using GoPro video cameras mounted inside the TOC on the wall opposite the screen.

We measured survival and search time to evaluate predator responses to moths. Each metric was evaluated at two time points: the end of the first searching bout, which was the initial uninterrupted period during which the bird actively searched for the moth after it had landed, and the end of the trial (i.e., after either two minutes of active searching or five minutes of total trial duration, whichever occurred first). To identify the first-bout of searching, we coded bird search behavior from trial videos using BORIS software (Friard and Gamba 2016). We coded the bird’s location (i.e., perch, screen, or floor) and head orientation (whether it was looking at or away from the screen) (see Supplementary Materials for details). Survival was binary; - moths that were not relocated at a given time point recorded as surviving. First-bout search time was the duration of the first search bout coded in the trial video, and whole trial search time was the total time delay between the moth landing and successful prey relocation and was also extracted from the trial videos.

The flying moths depicted in each trial were pre-rendered for computational efficiency. We generated 64 “flight sequences” that depicted flying moths (10 of each hindwing-background combination); the moth in each flight sequence had a unique flight trajectory, in which it flew between randomly selected points on the screen (see supplementary materials for more details on generating flight sequences). No bird experienced the same flight sequence more than once, and each sequence was presented to an average (median) of 6 birds (range 3-11) across the experiment. Each bird typically completed all 18 trials of the experiment within one day, and trials were run back-to-back unless the bird became unmotivated, in which case it was given a break (approx. 20 minutes).

### Landing Site Metrics

For each trial, the moth flew around the screen before landing at a random location and remaining in resting position with its hindwings concealed. Once landed, the moth blended into the background using its cryptic forewings, which fully covered the hindwings (Figure 2). Natural variation in background appearance, both between and within the six background images, together with differences between flight trajectories, could influence predator tracking and prey detectability. To account for this variation, we quantified both flight characteristics and camouflage-related properties of the stimuli.

For flight characteristics, we measured linear displacement as the distance (in pixels) between the moth’s initial position and its landing. Although all moths travelled the same flight distance, moths that landed closer to their starting position may have been easier for the birds to track and relocate after landing.

To account for differences in detectability of the landed moth, several camouflage metrics were calculated: edge disruption, pattern difference, luminance JND, and color JND. Prior to measurement, images of the moth in their landing position were converted to cone-catch images using the micaToolbox (Troscianko and Stevens 2015). These images use visual modeling to estimate how a given visual system would perceive a scene (Kelber et al. 2003; Endler and Mielke 2005). We generated cone-catch images using the blue tit (*Cyanistes caeruleus*) visual system (Hart et al. 2000), which was available in the micaToolbox (Troscianko and Stevens 2015). Because stimuli were displayed on RGB screens lacking ultraviolet output, the UV-sensitive channel was excluded from chromatic analyses, resulting in the use of an LMS rather than an LMSV visual model. For luminance, pattern, and edge disruption measures, we used the cone catch values of the double cone (dbl), which encodes luminance (achromatic brightness) perception in birds (Kelber et al. 2003; Endler and Mielke 2005). Edge disruption was calculated using GabRat, a measure which utilizes Gabor filters to calculate the ratio of false edges (at right angles to the outline) to coherent edges (parallel to the outline) (Troscianko et al. 2017); this metric quantifies the extent to which the moth could be cryptic in its landing site due to disruptive coloration (Cuthill et al. 2005). Pattern differences were calculated by measuring pattern energy (standard deviation, SD) at different spatial scales using Difference-of-Gaussian filters. Four spatial octaves were used: 1/8th, 1/16th, 1/32nd, and 1/64th of the moth width in pixels (121 px) (Troscianko et al. 2016; Barnett et al. 2021; Hancock et al. 2023). These measures were taken for both the moth and the immediate local background (circle of a diameter 2.5x the size of the moth), with the difference calculated as the Euclidean difference in energy for each scale between the moth and its background.

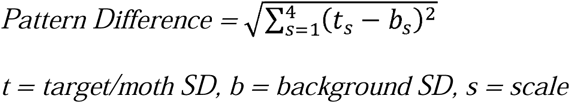

Pattern difference metrics quantify the similarity between two patterns, and therefore the similarity between the resting moth and the local bark background where it landed. Both the GabRat and pattern difference metrics used measurements from the blue tit dbl channel.

We used the receptor noise model to measure the distinctiveness between moths and their local backgrounds in just noticeable difference (JND) units, where one JND is the smallest color difference that can be reliably detected in good lighting conditions (Vorobyev and Osorio 1998; Siddiqi et al. 2004). We calculated luminance JNDs – which measure achromatic brightness differences – using the Siddiqi method based on a Weber fraction estimate for a budgerigar (v = 0.18; Lind et al. 2013) comparing the mean luminance of the moth forewings to that of its local background. Color JNDs – which measure chromatic differences independent of luminance – were calculated by converting the cone-catch images into RNL chromaticity images with the default weber fraction of 0.05 and blue tit cone ratios (sws = 1.92, mws = 2.68, lws = 2.70) (Hart et al. 2000), with color JND being equal to the Euclidean distance of the X and Y opponent color channels between the moth and its local background.

### Statistical Analysis

We analyzed digital moth survival probability and bird search times using mixed-effects models fit using the lme4 package (Bates et al. 2015) in R (v4.3.2; R Core Team, 2023). Survival probability was fit as the response variable in generalized linear mixed-effects models (GLMMs) with binomial error distributions and logit link functions. We conducted two analyses of moth survival (binary), using different metrics: first-bout survival, whether the prey survived the first predator searching bout, and trial survival, whether the prey survived the entire trial. Predator search time was fit as the response variable in linear mixed-effects models (LMMs). The search time analysis only included trials in which the moth was caught. Two search time metrics were analyzed separately: first-bout search time, defined as the period between when the moth lands and when the bird first looks away from the screen (with its head facing directly away from the screen), and whole-trial search time, defined as the period between when the moth lands and when it was pecked, irrespective of bird behavior. For both analyses search time was log-transformed to satisfy model assumptions.

Our analyses included the predictor variables hindwing color (categorical: red, yellow, cryptic), background (categorical: smooth, bark), season (categorical: spring, autumn), trial order, GabRat (numeric: edge disruption at landing location), pattern difference (numeric: Euclidean difference in pattern energy across spatial scales between moth and background at landing location), color JND (numeric: chromatic moth-background difference at landing location), luminance JND (numeric: achromatic moth-background difference at landing location), linear displacement (numeric: distance between spawn and land positions), and touchscreen chamber (TOC, categorical: 1, 2, or 3). Prior to model fitting, we assessed collinearity among continuous covariates, and excluded pattern difference from further analyses because it was moderately correlated with GabRat (r = 0.64). All continuous variables were rescaled to Standard Scores 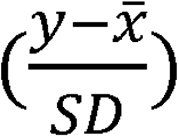 for use in statistical models.

For each model, we included the predictors above as fixed effects and individual bird ID (factor: unique number for each bird) and stimulus flight sequence (factor: unique number for each sequence) as crossed random intercepts. We did not include bird age as a fixed effect because the sample was small and unevenly distributed across age classes, limiting power to estimate age effects reliably. TOC and season were excluded from the whole-trial survival model because the restriction to bark-background trials resulted in unbalanced levels within both covariates and strong confounding between them, preventing their effects from being reliably estimated independently. The random-effects structure was simplified for both survival models, stimulus flight sequence was removed because the model was initially singular and its estimated variance approached zero. To improve model convergence, binomial survival models were fitted using the `bobyqa` optimizer. An interaction between color and background was initially included but removed as it was not significant. We evaluated the statistical significance of each predictor using likelihood ratio tests, implemented using the drop1 function in R. We assessed all model assumptions using the DHARMa package (Hartig 2022) in R. Simulated residuals were inspected for uniformity, dispersion, and zero-inflation, and residual plots were visually checked for any systematic patterns against fitted values or predictors.

## Results

### First-bout survival

First-bout survival probability was strongly affected by background, with trials on bark backgrounds resulting in higher survival rates than those on smooth backgrounds (χ²= 45.23, df= 1, p < 0.001; Fig. 4A). Linear displacement also impacted survival with shorter distances from the moth start and end location reducing survival (χ²= 10.12, df= 1, p = 0.001). Greater edge disruption (GabRat) increased moth survival (χ²= 4.41, df= 1, p = 0.036). Trial order didn’t significantly influence survival, but later trials tended to have lower moth survival (χ²= 3.32, df = 1, p = 0.069). In contrast, hindwing color did not significantly affect first-bout survival (χ² = 3.45, df = 2, p = 0.178). Neither did season (χ²= 0.02, df= 1, p = 0.886), color JND (χ²= 3.10, df= 1, p = 0.078), TOC (χ²= 0.73 df= 2, p = 0.695), or luminance JND (χ²= 0.002 df= 1, p = 0.961),

**Figure 4.**
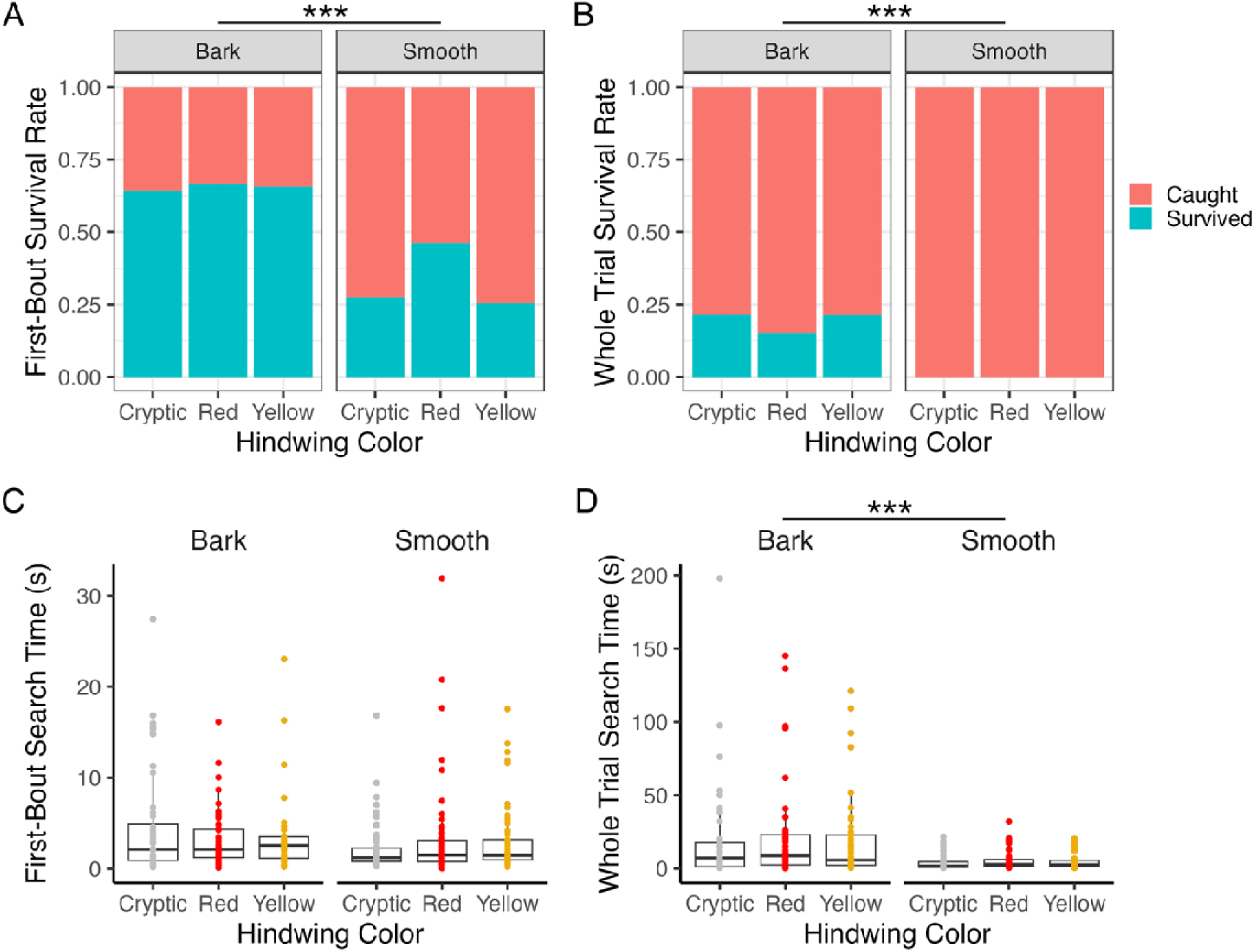
Effect of hindwing color on digital moth survival and search time on bark and smooth backgrounds. (A) Survival rate during the first bout of continuous predator searching. (B) Survival rate for the whole trial. (C) Search time for the first bout. (D) Search time for the whole trial. Hindwing colors are cryptic, red, and yellow. Asterisks (***) indicate a statistically significant difference between backgrounds (*p* < 0.001). No significant differences were detected among hindwing color treatments within either background.

### Whole trial survival

Whole trial survival differed strongly between backgrounds, with no moths surviving to the end of the trial on smooth backgrounds (Fisher’s exact test: p < 0.001; Fig. 4B). Consequently, analyses of whole-trial survival were restricted to bark-background trials only. Within bark trials, luminance contrast between the moth and background had a strong effect on survival, with higher luminance JNDs associated with a reduced probability of surviving the entire trial (χ² = 10.796, df = 1, p =0.001). In contrast, whole-trial survival probability was not affected by hindwing color (χ² = 0.401, df = 2, p = 0.818). No other variables significantly affected whole trial survival (See supplementary results).

### First-bout search time

First-bout search time was influenced by trial order, with predators becoming faster at locating prey over successive trials, consistent with learning or increased search efficiency (χ² = 5.283, df = 1, p =0.022; Fig. 4C). In contrast, hindwing color did not significantly affect first-bout search time (χ² = 0.772, df = 2, p =0.680), and there were no significant effects of background (χ² = 1.252, df = 1, p =0.263) or season (χ² = 0.905, df = 1, p =0.341). None of the landing site metrics, linear displacement or TOC significantly predicted first-bout search time (see supplementary results).

### Whole trial search time

Whole trial search time differed strongly between backgrounds, with moths presented on smooth backgrounds found more rapidly than those on bark backgrounds (χ² = 27.957, df = 1, p <0.001; Fig. 4D). Linear displacement also significantly influenced total search time, with moths landing farther from their initial position taking longer for predators to locate (χ² = 10.638, df = 1, p =0.001). In contrast, hindwing color did not significantly affect whole trial search time (χ² =1.012, df = 2, p =0.603), and there were no significant effects of any other covariates (see supplementary results).

## Discussion

We found no evidence that conspicuous hindwing coloration of digital *Catocala*-like moths increased prey survival or impaired the ability of predators to relocate prey. Wild blue tits relocated moths with conspicuous and cryptic hindwings equally effectively across both survival and search-time metrics. However, conspicuous hindwings also caused no detectable predation cost relative to cryptic hindwings. In contrast, predator responses were strongly influenced by background type, landing site measure (local luminance, edge disruption) flight displacement, and repeated trial exposure, demonstrating that the touchscreen experimental setup successfully captured biologically meaningful variation in predator search behavior.

One explanation for the absence of a detectable survival advantage of flash coloration is that any advantage conferred by conspicuous hindwing displays may be subtle, transient, or highly context dependent. Flash coloration has been hypothesized to disrupt predator tracking or prey relocation immediately following escape, yet such effects may depend strongly on factors such as flight speed, escape trajectory, viewing distance, background structure, or predator experience. Several studies suggest that protective coloration can be more effective against naïve predators than experienced ones (Lyytinen et al. 2004; Umbers et al. 2019).

Across repeated trials, predators rapidly improved their search efficiency within the first bout of continuous searching, suggesting that any transient confusion caused by conspicuous hindwing displays may have been quickly overcome through learning, but these learning effects may be less pronounced in more complex ecological contexts. Similarly, because moths remained continuously visible during flight on the touchscreen, predators may have retained accurate information about prey landing position regardless of hindwing coloration, although search time and survival still increased for bark backgrounds. Under natural conditions, prey movement through more visually complex environments may create greater uncertainty about prey relocation following landing. In particular, the more complex three-dimensional nature of predator-prey interactions in natural habitats may be more favorable for any antipredator benefits of flash coloration than simulations on two-dimensional screens.

Viewed together with previous experimental studies (Table 1), our results suggest that the anti-predator effects of flash coloration are neither universally strong nor consistently expressed across systems. Although several studies have reported survival advantages associated with movement-dependent conspicuous signals, others — including our study — have found weak or absent effects. Importantly, studies differ substantially in predator identity, prey realism, movement dynamics, and the ecological context in which predator decisions are made. Collectively, this emerging pattern suggests that flash coloration may not represent a single generalized anti-predator mechanism, but instead a context-dependent phenomenon whose effectiveness depends on the interaction between prey movement, predator perception, and environmental structure.

While flash coloration did not provide any antipredator benefits in our experiments, it also did not have any predation costs, i.e. a reduced survival because the conspicuous color was easier to follow. This suggests that flash coloration may result from relaxed selection, or could represent a trade-off between predator avoidance and another selective function, such as sexual selection. This would effectively make flash coloration an extreme version of signal partitioning, in which conspicuous signals are restricted to body regions that are more visible to receivers (often conspecifics) than to predators (Marshall and Stevens 2014). To the extent that flash coloration allows the evolution of conspicuous coloration on at least some body regions without the typical predation costs, it may also be a relatively inexpensive pathway for the evolution of conspicuous coloration more broadly, including warning and mating signals (Loeffer-Henry et al. 2025). Although *Catocala* moths are widely regarded as classic examples of putative flash coloration, conspicuous hindwing displays may also serve alternative or additional functions. For example, sudden exposure of brightly colored hindwings could contribute to a startle effect (deimatic display) during close-range predator encounters (Kang et al. 2017; Umbers et al. 2017; Kim et al. 2020). However, such mechanisms typically depend on rapid prey display at short distances and may differ behaviorally from the relocation-based mechanism proposed under the flash coloration hypothesis.

Our results provide no strong support for the flash coloration hypothesis in *Catocala*-like prey under the conditions tested, suggesting that conspicuous hindwing displays alone may not consistently disrupt predator tracking or prey relocation. However, the absence of detectable predation costs, together with the inconsistent effects reported across previous studies, suggests that any benefits of flash coloration are likely to be conditional rather than universally expressed. More broadly, our results demonstrate the value of touchscreen-equipped operant chambers for studying dynamic predator-prey interactions using wild predators under controlled conditions. Our experiment generated biologically meaningful predator responses to moving prey, reinforcing that touchscreen systems can provide a powerful framework for experimentally testing hypotheses involving dynamic visual signals that are otherwise difficult to manipulate consistently using live prey or analog prey models. Expanding touchscreen research beyond lab animals and humans bridges the gap between tightly controlled lab experiments and ecologically relevant studies. It allows research into how wild animals perceive and respond to complex visual signals while retaining high experimental control of stimuli.

## Acknowledgements

We would like to thank the Research Council of Finland (grant numbers 355955 to JM) for funding this research, Lammi Biological Station where this research took place and, the EEI research group, James Barnett and Andy Hurly for the valuable feedback and discussion. We are especially grateful to Cristina Jusdado, Emilie Favaro and Fiorina Muster for assisting with the data collection.

## Permits

Wild birds were used with permission from the Central Finland Regional Environment Centre (VARELY/6870/2022).

## Data Availability Statement

The supporting data will be archived in an appropriate public repository, and the data DOI will be included upon acceptance.

