## Supplementary Material for "Testing The Protective Function of Flash Coloration Using Wild Avian Predators"

### Supplementary materials

***Supplementary Methods***

*Training Procedure*

Pre-training: To train birds to positively associate pecking the moth image with receiving a reward, birds were first trained to peck paper moths. To prevent any bias towards either experimental color these paper moths were colored either light red (the midpoint of the test red color and the test grey color) or light yellow (the midpoint of the test yellow color and the test grey color). Initially birds were presented with an array of three light yellow and three light red paper moths on a piece of cardboard with a half-sunflower seed glued loosely on each one. Once the bird had removed all rewards, they were presented with a second array where the half seeds were hidden under wings and within the body of the moths. The aim of this second array was to motivate the bird to attack the moths and not just the food reward. The final stage of the pre-training was familiarization with the reward system (described below) where the reward box, fixed open and filled with sunflower seeds, replaced the normal feeding tray until all seeds were eaten. This allowed birds to overcome the initial neophobia associated with the reward box.

Paper-training: Birds were presented with a single paper moth close to the reward box. The reward box consisted of a small remote-controlled servo motor attached to an arm that could raise and lower the lid to an opaque plastic box filled with food. The mechanism of the box was hidden inside a thin plastic box cut for the purpose and covered in brown paper to prevent the motor and arm from being visible. Once the bird had successfully pecked the moth or showed interest in it (e.g., staring at or standing on the stimuli), the reward box was opened. The appearance of the moth randomly varied between the light-red and light-yellow morph, such that both morphs were presented an equal number of times across every six presentations. As the bird’s moth-directed behaviors increased in frequency, only successful pecks of the paper moth were rewarded, and this process was repeated until the bird was regularly pecking a moth and then taking a seed. The location of the paper moth was then moved farther from the reward box on the floor of the TOC and then onto the screen. This continued until the researcher was satisfied that the bird could confidently peck the paper moth and then take a reward. Breaks were given to the birds whenever they showed a lack of motivation.

Digital training: The digital training was conducted on the touchscreen setup (the touchscreen-equipped operant chamber (TOC) is described in the main text). MATLAB (v. R2022B) was used to create the touchscreen digital training game. The digital training was broken into 6 steps; each training step was repeated until the researcher was satisfied that the bird was confident pecking stimuli anywhere on the screen (with a minimum of 5 trials) for that step. Between every training trial (each unique presentation within a training step), a grey screen was displayed, and the trial was started only when the bird was on the perch and facing the screen. Between each training step, the monitor was turned off and the screen was dark. Training steps were as follows:

1. A grey screen was displayed with a digital moth modeled on a *Catocala* moth at rest (hindwings covered), the bird had to peck the moth, the moth would disappear, and then the reward box was opened to allow the bird to take a reward. As with the paper training, these moths were colored light red or light yellow with the same criteria for equal presentations. There was a buffer zone of 15 pixels around the moth where the game would accept a peck on the moth to allow for small errors in touch frame accuracy. For the training, the moth had two possible sizes, either large (7172 total pixels; 3 x 2.5cm) or small (2879 pixels; 2 x 1cm). The smaller size was used as initially we planned to carry out this experiment with two moth sizes.
2. A grey screen with a digital resting moth was displayed. Once the bird pecked the moth, it would slowly move 100 pixels (~3cm) in approximately 2.5 seconds and the bird had to re-peck the moth before receiving a reward. For this step, the resting moth simply glided while moving - there was no flapping animation. This step was needed because during the experiment, birds had to be trained to peck the moth twice, once to initiate the flight of the moth and again to make the moth disappear before receiving a reward. This meant when the bird started an experimental trial, it was motivated to find the moth after it landed (i.e., after the second peck). With all moving stimuli, the moth had to ‘land’ (stop moving) before the screen would register a peck on the stimulus as a ‘hit’, yielding a reward and the conclusion of the trial.
3. As in step two, but the moth glided 500 pixels (~14cm) in approximately 2.5 seconds between pecks.
4. As in step three, but within each trial after the first peck on the moth the grey background was replaced with a low contrast bark background (see below, “Background images”) behind the moving moth. The background bark image was randomly selected from 3 potential images for each training trial. This training step was included to gradually introduce the birds to the textured backgrounds, which some were initially hesitant to interact with. In the low contrast background images, the background is less salient and the moth is still easily distinguishable from the background.
5. As in step four, but with a medium contrast version of the bark background image.
6. As in step five, but with a full contrast version of the bark background image.

For each trial, bird ID, trial ID, peck time, peck coordinates, whether the peck was on the stimuli (hit) or not (miss), and the start point and landing point of the moth were recorded automatically by MATLAB. A number of birds (8) that started the digital training did not progress through all steps and so were released and excluded from the experiment.

*Generating stimuli*

Moth stimuli: The moth stimuli used in the tests were based on *Catocala* *concumbens*. A pinned *C. concumbens* specimen from Luonnontieteellinen keskusmuseo (Finnish Museum of Natural History) was photographed in RAW format using a Sony A6000 camera under a Powersun H.I.D Metal Halide reptile lamp, with a color temperature of 6500K, alongside a color standard and a ruler. The image was converted to a 16-bit linear tiff, standardized for color using an Avian Technologies 50% reflectance grey standard, and scaled to 3 pixels per mm. The moth was then segmented from the background, and rotated such that the two points where the outer edges of the forewings and hindwings met formed a horizontal line. Within the moth, five regions of interest (ROIs) were then marked: left forewing, right forewing, left hindwing, right hindwing, and body (see Figure S1; ROIs outlined in green). For each wing ROI, a ‘hinge point’ was marked as the point at which the wing intersected with the body (see Figure S1; hinge points marked in blue). For each hindwing, the colorful portion of the wing was segmented from the melanized portion. To generate a moth in resting posture from the spread-wing configuration, forewing ROIs were rotated by 90° and hindwing ROIs by 45° in the direction of natural wing folding (i.e. left wings rotated anticlockwise, right wings clockwise), such that the forewings fully covered the hindwings.

Using the processed *C. concumbens* image, we replaced the colored portions of the hindwings to create three experimental stimuli: (1) the cryptic stimulus with hindwings colored grey (RGB (uint8): 125, 117, 113), calculated as the mean RGB of the forewings; (2) conspicuous red (RGB: 248, 42, 42) and (3) conspicuous yellow (RGB: 217, 144, 16) stimuli with hindwing colors that approximated the most intense red and yellow hindwings observed in extant moths in the genus *Catocala*.

Training stimuli were generated by replacing the forewings with a solid color (i.e., training stimuli had no internal patterning on the wings), and rotating the wings into the resting position, with the forewings covering the hindwings. Forewings were rotated down 90 degrees around their hinge points and hindwings were rotated up 45 degrees around their hinge points. To prevent generating color biases at the training stage, we used greyish-red (RGB: 196, 88, 86) and greyish-yellow (RGB: 177, 131, 81) forewing colors for the training stimuli, with each training color calculated as the midpoint between the grey hindwing color and the red and yellow hindwing colors, respectively.

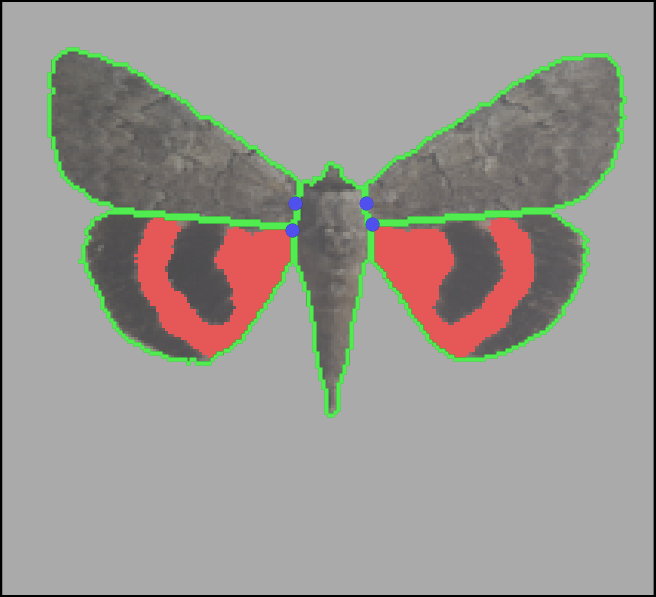

**Figure S1**. Definition of regions of interest (ROIs) and wing hinge points used to generate moth flight animations. The moth is shown in the spread-wing configuration with five ROIs outlined in green: the body, left and right forewings, and left and right hindwings. Blue points indicate the hinge positions for each wing, defined as the midpoint of the wing–body attachment. These hinge points were used as fixed points when rotating the wings to simulate wing opening, closing, and scissoring motion during flight.

When moths were presented in the resting position at the beginning of trials, hindwing ROIs were removed to prevent them from being visible at the wing margins. All flight sequences began from this resting configuration. The moth flight animation was generated by rotating the wings in a scissoring motion.

We generated flying moths from static stimuli by systematically rotating wing ROIs around their respective hinge points. Flight began with a wing-opening sequence, during which the wings rotated dorsally at 15° per frame until reaching the fully spread position. During the first four frames, only the forewings moved, followed by coordinated movement of both forewings and hindwings. The opening sequence consisted of the following angular rotations (in degrees) across successive frames: left forewing [90, 75, 60, 45, 30, 15, 0], right forewing [-90, -75, -60, -45, -30, -15, 0], left hindwing [45, 45, 45, 45, 30, 15, 0], and right hindwing [-45, -45, -45, -45, -30, -15, 0]. Once fully extended, wing movement followed a repeating scissoring cycle at 12° per frame, in which forewings and hindwings moved in opposition, such that the forewings repeatedly covered and revealed the hindwings. This cycle consisted of the following angular sequence: left forewing [0, 12, 24, 12], right forewing [0, -12, -24, -12], left hindwing [0, -12, -24, -12], and right hindwing [0, 12, 24, 12]. This sequence was repeated until landing. Landing involved a wing-closing sequence that was the inverse of the opening sequence. In experiment trials, the entire moth configuration (body and wings at the appropriate phase of the flight cycle) was translated across the screen to simulate flight.

Background images: Moth stimuli were presented against background images that depicted either tree bark or were solid grey. Tree bark images were created based on images of six willow (*Salix sp.*) trees in Tartu, Estonia. Trees were selected opportunistically, and photographed under ambient lighting using the same equipment as moth photography. Tree images were standardized and scaled using the same procedure described above, then cropped to the region containing predominantly forward-facing tree bark (i.e., excluding tree edges to avoid aberrant texture and lighting effects). Because the trees were not wide enough to cover the whole TOC screen at life-size resolution, we tiled the images horizontally to create the appearance of solid tree bark image across the whole screen. Three tree bark images were used as experimental backgrounds. Each of these tree bark images was paired with a solid grey version, which was the mean color of that bark image. Three additional tree bark images were used as training backgrounds; for each of these images, we created low contrast and medium contrast versions, in addition to the original (full contrast) image. Low contrast images were created by calculating the mean of the bark image and the respective solid grey image with a weight of [1,3] (i.e., 25% bark image, 75% grey image) for each pixel; medium contrast images were created by calculating the unweighted mean of the bark image and the respective solid grey image (i.e., 50% bark image, 50% grey image).

All stimuli were presented after adding a square root transformation to linear images, to mimic nonlinear visual perception. All color-based calculations were done based on measurements from linear images. RGB values reported above are the colors shown (i.e., post-transformation).

*Trial Procedure*

Before starting a trial, a grey screen (RGB: 0.71, 0.71, 0.71) was displayed until the bird appeared ready to start the trial, generally this meant it was on the perch, looking motivated, and not resting, bathing, or drinking water. The initial screen was then displayed showing the background image covering most (~70%) of the screen, bordered by a 6cm grey (RGB: 0.71, 0.71, 0.71) frame. The moth in a resting position was randomly positioned in this grey border as follows: first the spawn side – top, bottom, left, or right – was randomly chosen, then a random point at least 120 pixels from the outer edges of that side was chosen and the resting moth was inserted with the middle of the body on this point; the moth was always equidistant between the inner and outer boundaries of the frame (so always the same distance from the edge of the monitor), and resting in a vertical position (i.e., head up). The background on the initial screen was same as would be used in the trial, either a bark or smooth image. The moth was displayed on the grey border to make it easy to see and therefore easy for the bird to peck to start the trial. The trial started when the bird pecked the moth (or within a 15 pixel buffer zone around the edges of the moth), which caused the moth to initiate flight. Having the birds initiate the trials via the first peck ensured that they were paying attention and motivated to find the moth after it landed to gain a food reward. Occasionally, the infrared touch frame would not register a peck, in which case a manual override was triggered by the researcher to generate a peck. Immediately after being pecked, the moth would start its 2.3 second flight. On the first frame of the flight sequence, the grey border was removed so that background image covered the entire screen. The moth flapped its wings, displaying either cryptic, red, or yellow hindwings. Moth flight was simulated as a series of straight flight segments between randomly selected target points on the screen. At each step, the stimulus selected a new location and flew directly to it at a constant speed before selecting the next target. The range of possible coordinates was selected so the moth was always fully visible on the screen. The moth was always orientated with its head up throughout its movement. After 2.3 seconds, the moth stopped and landed – its forewings returned to the resting position and the hindwings were hidden. When the moth landed, the observer started a stopwatch, recording the active search time of the bird. The timer was stopped whenever the bird was not looking at the screen and restarted when the bird turned back towards the screen. Once the moth landed, the bird had to find the moth on the screen and peck it (or within the 15 pixel buffer). If the bird pecked the moth, the screen immediately returned to a grey (RGB=0.71, 0.71, 0.71) screen and the bird was rewarded. The reward box was manually operated by the observer from outside the TOC using a remote control. The bird was allowed to take one half sunflower seed from the box. Occasionally the bird would remove more than one half seed, in which case the extra seed was removed. If the bird did not peck the moth within either two minutes of active searching or five minutes of total trial time, the trial was ended and the bird did not get a reward. See Supplementary Video 1 for a trial video.

*TOC screen settings*

One TOC screen was found to be set to 60 Hz, rather than the intended 165 Hz, partway through the trial. It is possible that this may have affected how the birds viewed this screen, although we note that these are flicker-free screens so even at this lower framerate they would not have appeared to be flickering on and off. We included TOC identity as a variable in our models to account for any equipment differences; this variable was never a statistically significant predictor, suggesting such effects were minimal. We also note that all moving stimuli in our games were rendered at 60 frames per second, which is below the critical flicker fusion threshold of blue tits (Boström et al. 2016). Stimulus movement may therefore have been perceived as jumpy to the bird subjects due to lower frame rate, although any movement effects would have been consistent across all trial conditions and are therefore unlikely to influence our general pattern of results.

*Video coding in BORIS*

We documented the behavioral responses of birds to digital moth stimuli by coding videos of each trial. We coded two state behaviors: the bird’s location and head orientation. The bird’s location was coded as its position in the TOC, with the options perch (feet in contact with the perch, with the body either positioned above or hanging below), screen (feet in contact with the wire grid that facilitated access to the experimental screen, or jumping between wires on the grid or jumping to peck the stimuli), or floor (standing anywhere on the floor of the TOC). Other locations were not recorded, nor was time spent flying between locations. The bird’s head orientation could be either looking at or away from the screen. A bird was classified at looking at the screen unless the head was facing directly away from the screen or if it was preening or drinking water (Tyrrell and Fernández-Juricic 2017). This yielded a conservative estimate of time spent looking at the screen, since birds were classified as looking at the screen if there was any possibility that they may have been able to see the screen. For each state behavior, we recorded the beginning and end of the transition to that state, resulting in a duration for each bout of the behavior. The first search bout therefore began when the bird first oriented its head toward the screen after the moth had landed. The bout continued while the bird’s head orientation indicated that the screen could fall within its visual field, and ended when the bird’s head orientation was classified as looking directly away from the screen.

***Supplementary Material References***

Boström JE et al. 2016. Ultra-Rapid Vision in Birds Neuhauss SCF, editor. PLoS ONE. 11(3):e0151099. https://doi.org/10.1371/journal.pone.0151099

Tyrrell LP, Fernández-Juricic E. 2017. Avian binocular vision: It’s not just about what birds can see, it’s also about what they can’t Osorio D, editor. PLoS ONE. 12(3):e0173235. https://doi.org/10.1371/journal.pone.0173235

***Supplementary Results***

*Models*

Model 1. First-bout survival – LRT table

| Predictor | χ² | df | p-value |
| --- | --- | --- | --- |
| Color | 3.45 | 2 | 0.178 |
| Background | 45.23 | 1 | 1.75 e-11 |
| Season | 0.02 | 1 | 0.886 |
| Trial_order | 3.32 | 1 | 0.069 |
| GabRat (scaled) | 4.41 | 1 | 0.036 |
| Color_JND (scaled) | 3.10 | 1 | 0.078 |
| Luminance_JND (scaled) | 0.00 | 1 | 0.961 |
| Displacement (scaled) | 10.12 | 1 | 0.001 |
| TOC | 0.73 | 2 | 0.695 |

Model 1. First-bout survival – Summary table

| Predictor | Estimate | Standard Error |
| --- | --- | --- |
| Intercept | 1.666 | 0.583 |
| Color_red | 0.353 | 0.335 |
| Color_yellow | -0.216 | 0.328 |
| Background_smooth | -2.391 | 0.401 |
| Season_2 | -0.096 | 0.669 |
| Trial_order | -0.043 | 0.024 |
| GatRat_scaled | 0.402 | 0.199 |
| Color_JND (scaled) | -0.237 | 0.135 |
| Luminance_JND (scaled) | -0.007 | 0.137 |
| Displacement (scaled) | 0.457 | 0.147 |
| TOC_2 | 0.073 | 0.672 |
| TOC_3 | -0.520 | 0.752 |

Intercept: Color_cryptic, Background_bark, Season_1, TOC_1

Model 2. Whole trial survival – LRT table

| Predictor | χ² | df | p-value |
| --- | --- | --- | --- |
| Color | 0.401 | 2 | 0.818 |
| Trial_order | 0.121 | 1 | 0.728 |
| GabRat (scaled) | 3.008 | 1 | 0.083 |
| Color_JND (scaled) | 0.392 | 1 | 0.531 |
| Luminance_JND (scaled) | 10.796 | 1 | 0.001 |
| Displacement (scaled) | 0.650 | 1 | 0.420 |

Model 2. Whole trial survival – Summary table

| Predictor | Estimate | Standard Error |
| --- | --- | --- |
| Intercept | -2.556 | 0.969 |
| Color_red | 0.313 | 0.705 |
| Color_yellow | -0.146 | 0.649 |
| Trial_order | -0.017 | 0.048 |
| GabRat (scaled) | 0.456 | 0.265 |
| Color_JND (scaled) | 0.208 | 0.331 |
| Luminance_JND (scaled) | -1.270 | 0.445 |
| Displacement (scaled) | -0.274 | 0.343 |

Intercept: Color_cryptic

Model 3. First-bout search time – LRT table

| Predictor | χ² | Df | p-value |
| --- | --- | --- | --- |
| Color | 0.772 | 2 | 0.680 |
| Background | 1.252 | 1 | 0.263 |
| Season | 0.905 | 1 | 0.341 |
| Trial_order | 5.283 | 1 | 0.022 |
| GabRat (scaled) | 0.091 | 1 | 0.762 |
| Color_JND | 0.042 | 1 | 0.838 |
| Luminance_JND (scaled) | 0.019 | 1 | 0.890 |
| Displacement (scaled) | 1.571 | 1 | 0.210 |
| TOC | 2.229 | 2 | 0.328 |

Model 3. First-bout search time – Summary table

| Predictor | Estimate | SE | Corrected Estimate | Corrected SE |
| --- | --- | --- | --- | --- |
| Intercept | 0.886 | 0.237 | 2.426 | 0.575 |
| Color_red | -0.085 | 0.173 | 0.919 | 0.159 |
| Color_yellow | 0.035 | 0.171 | 1.036 | 0.177 |
| Background_smooth | -0.213 | 0.201 | 0.808 | 0.162 |
| Season_2 | 0.206 | 0.244 | 1.229 | 0.300 |
| Trial_order | -0.024 | 0.011 | 0.976 | 0.011 |
| GabRat (scaled) | -0.032 | 0.101 | 0.969 | 0.098 |
| Color_JND (scaled) | 0.009 | 0.069 | 1.009 | 0.070 |
| Luminance_JND (scaled) | 0.003 | 0.071 | 1.003 | 0.071 |
| Displacement (scaled) | 0.088 | 0.074 | 1.092 | 0.081 |
| TOC_2 | -0.034 | 0.247 | 0.967 | 0.239 |
| TOC_3 | -0.350 | 0.275 | 0.704 | 0.193 |

Intercept: Color_cryptic, Background_bark, Season_1, TOC_1

For corrected estimates, the intercept represents the first-bout search time in seconds and the co-variate estimates represent proportions. The SE were estimated by multiplying the SE by the corrected estimated.

Model 4. Whole trial search time – LRT table

| Predictor | χ² | Df | p-value |
| --- | --- | --- | --- |
| Color | 1.012 | 2 | 0.603 |
| Background | 27.957 | 1 | 0.000 |
| Season | 0.538 | 1 | 0.463 |
| Trial_order | 2.562 | 1 | 0.109 |
| GabRat (scaled) | 1.588 | 1 | 0.208 |
| Color_JND | 0.106 | 1 | 0.744 |
| Luminance_JND (scaled) | 0.246 | 1 | 0.620 |
| Displacement (scaled) | 10.638 | 1 | 0.001 |
| TOC | 2.931 | 2 | 0.231 |

Model 4. Whole trial search time – Summary table

| Predictor | Estimate | Standard Error | Corrected Estimate | Corrected SE |
| --- | --- | --- | --- | --- |
| Intercept | 2.203 | 0.249 | 9.050 | 2.255 |
| Color_red | 0.185 | 0.200 | 1.204 | 0.241 |
| Color_yellow | 0.070 | 0.197 | 1.072 | 0.212 |
| Background_smooth | -1.265 | 0.234 | 0.282 | 0.066 |
| Season_2 | 0.144 | 0.220 | 1.154 | 0.253 |
| Trial_order | -0.021 | 0.013 | 0.979 | 0.013 |
| GabRat (scaled) | 0.145 | 0.119 | 1.156 | 0.137 |
| Color_JND (scaled) | -0.030 | 0.079 | 0.970 | 0.077 |
| Luminance_JND | 0.033 | 0.081 | 1.034 | 0.084 |
| Displacement (scaled) | 0.273 | 0.086 | 1.314 | 0.113 |
| TOC_2 | -0.051 | 0.224 | 0.950 | 0.213 |
| TOC_3 | -0.367 | 0.250 | 0.693 | 0.173 |

Intercept: Color_cryptic, Background_bark, Season_1, TOC_1

For corrected estimates, the intercept represents the whole bout search time in seconds and the co-variate estimates represent proportions. The corrected SEs were estimated by multiplying the SE by the corrected estimated.
